# PDGFR dimer-specific activation, trafficking and downstream signaling dynamics

**DOI:** 10.1101/2021.10.26.465978

**Authors:** Madison A. Rogers, Katherine A. Fantauzzo

**Affiliations:** Department of Craniofacial Biology, School of Dental Medicine, University of Colorado Anschutz Medical Campus, Aurora, CO

**Author notes:** Correspondence to Katherine A. Fantauzzo.

## Abstract

Signaling through the platelet-derived growth factor receptors (PDGFRs) plays a critical role in multiple cellular processes during development. The two PDGFRs, PDGFRα and PDGFRβ, dimerize to form homodimers and/or heterodimers. Here, we overcome previous limitations in studying PDGFR dimer-specific dynamics by generating cell lines stably expressing C-terminal fusions of each PDGFR with bimolecular fluorescence complementation (BiFC) fragments corresponding to the N-terminal or C-terminal regions of the Venus fluorescent protein. We find that PDGFRβ receptors homodimerize more quickly than PDGFRα receptors in response to PDGF ligand, with increased levels of autophosphorylation. Further, we demonstrate that PDGFRα homodimers are trafficked and degraded more quickly, while PDGFRβ homodimers are more likely to be recycled back to the cell membrane. We show that PDGFRβ homodimer activation results in a greater amplitude of phospho-ERK1/2 and phospho-AKT signaling, as well as increased proliferation and migration. Collectively, our findings provide significant insight into how biological specificity is introduced to generate unique responses downstream of PDGFR engagement.

**Summary:** The authors utilize a novel bimolecular fluorescence complementation approach to investigate PDGFR homodimer-specific dynamics. They uncover differences in the timing and extent of receptor dimerization, activation and trafficking, which lead to changes in downstream signaling and cellular activity.

## Introduction

The platelet-derived growth factor receptors (PDGFRs) are a family of receptor tyrosine kinases (RTKs) known to direct multiple cellular processes during development, such as migration, proliferation and differentiation (Heldin and Westermark, 1999; Hoch and Soriano, 2003). In mammals, this family is composed of four dimeric growth factor ligands – PDGF-AA, PDGF-BB, PDGF-CC and PDGF-DD – which variously signal through two receptors, PDGFRα and PDGFRβ (Bostrom et al., 1996; Ding et al., 2004; Leveen et al., 1994; Soriano, 1994; Soriano, 1997; Williams, 1989). Each receptor consists of an extracellular region harboring five immunoglobulin-like loops, a single transmembrane domain and an intracellular domain containing a split, catalytic tyrosine kinase (Williams, 1989). Ligand binding results in the dimerization of two PDGFRs to form PDGFRα homodimers, PDGFRα/β heterodimers or PDGFRβ homodimers (Fantauzzo and Soriano, 2016; Herren et al., 1993; Matsui et al., 1989; Seifert et al., 1989; Shim et al., 2010), followed by activation of the tyrosine kinase domains and trans-autophosphorylation of intracellular tyrosine residues (Herren et al., 1993; Kelly et al., 1991). Signaling molecules possessing Src homology 2 phosphotyrosine recognition motifs subsequently bind to phosphorylated residues in the intracellular domains of the receptors and initiate downstream signaling cascades (Heldin and Westermark, 1999; Williams, 1989). When expressed in the same cell type, signaling through the two receptors can result in different cell activities. For example, in the mouse cranial neural crest cell lineage, PDGFRα plays a predominant role in cranial neural crest cell migration, while PDGFRβ primarily contributes to proliferation of the cranial neural crest-derived facial mesenchyme (Mo et al., 2020). Given the structural similarities of PDGFRs and the fact that they interact with similar subsets of signaling molecules (Heldin and Westermark, 1999), a critical question in the field is how this biological specificity is introduced to generate unique responses downstream of receptor engagement.

PDGFR internalization into the cell following ligand binding and dimerization also contributes to the regulation of PDGFR signaling (Bonifacino and Traub, 2003). Once internalized, PDGFRs can continue to bind adaptor proteins and/or signaling molecules within endosomes to activate downstream signaling pathways (Jastrzebski et al., 2017; Teis et al., 2002; Wang et al., 2004). Endosomal trafficking of the receptors commonly results in receptor degradation (Hellberg et al., 2009; Sadowski et al., 2013). However, two studies have demonstrated that PDGFRβ homodimers and PDGFRα/β heterodimers can also be recycled to the cell membrane (Hellberg et al., 2009; Karlsson et al., 2006), indicating potential dimer-specific differences in PDGFR trafficking following activation. Despite these findings, the dimer-specific dynamics of PDGFR activation, trafficking and signal attenuation, as well as the roles of these processes in mediating signal transduction and cellular activity, have not been studied in detail.

Here, we implemented bimolecular fluorescence complementation (BiFC) (Hu and Kerppola, 2003; Magliery et al., 2005) to probe for PDGFR dimer-specific dynamics. The BiFC technique adapted for this study employs a split Venus fluorescent protein fused to the C-terminus of individual PDGFRs to investigate receptor dimerization and has been used to investigate homodimeric and heterodimeric interactions of the RTK ERBB2 (Croucher et al., 2016; Kennedy et al., 2019). This approach overcomes limitations with prior approaches to examine receptor expression and/or dimerization using antibody-based methods that cannot distinguish between PDGFRs present as monomers or engaged in homodimers versus heterodimers. For this study, we generated two stable cell lines – one for each PDGFR homodimer pair – expressing BiFC-tagged PDGFRs. The formation of a functional Venus fluorescent protein upon PDGFR dimerization allows for the visualization and purification of individual PDGFR dimers. Utilizing this technique, we demonstrated significant differences in the timing and extent of PDGFRα versus PDGFRβ homodimer dimerization, activation and trafficking, which lead to changes in downstream signaling and cellular activity. Taken together, these data shed considerable light on the mechanisms by which biological specificity is introduced downstream of PDGFR activation.

## Methods and materials

### Generation of PDGFR-BiFC-HCC15 cell lines

Using Gateway cloning, we cloned pDONOR223-PDGFRA (Addgene 23892) and pDONOR223-PDGFRB (Addgene 23893) each into pDEST-ORF-V1 (Addgene 73637) and, separately, into pDEST-ORF-V2 (Addgene 73638). PDGFRα-V1, PDGFRα-V2, PDGFRβ-V1 and PDGFRβ-V2 sequences were amplified by PCR using the following primers:

5’-TTACTCGAGATGGGGACTTCCCATCCGGC-3’ and

5’-TTACCCGGGTTACTGCTTGTCGGCGGTGA-3’,

5’-TTACTCGAGATGGGGACTTCCCATCCGGC-3’ and

5’-TTACCCGGGTTACTTGTACAGCTCGTCCA-3’,

5’-TTAGAATTCATGCGGCTTCCGGGTGCGAT-3’ and

5’-CTGTCTAGATTACTGCTTGTCGGCGGTGA-3’,

5’-TTAGAATTCATGCGGCTTCCGGGTGCGAT-3’ and

5’-CGGTCTAGATTACTTGTACAGCTCGTCCA-3’, respectively. The sequences were cloned into the pLVX-Puro vector using XhoI and XmaI sites for PDGFRα-V1 and PDGFRα-V2 and EcoRI and XbaI sites for PDGFRβ-V1 and PDGFRβ-V2. The above lentiviral constructs (10 mg) and packaging vectors pCMV-VSV-G (Stewart et al., 2003) and pCMV-dR8.91 (Zufferey et al., 1997) (5 mg each) were transfected into HEK 293T/17 cells using Lipofectamine LTX (Thermo Fisher Scientific). Media containing lentivirus was collected 48 h and 72 h following transfection and filtered using a 13 mm syringe filter with a 0.45 µm PVDF membrane (Thermo Fisher Scientific) following addition of 4 mg/mL polybrene (Sigma Aldrich). Lentiviral-containing media was added to HCC15 cells every 24 h for two days, and cells were subsequently grown in the presence of 2 µg/mL puromycin for 10 days. Individual Venus-positive cells were isolated on a Moflo XDP 100 cell sorter (Beckman Coulter, Inc.) following 5 min of stimulation with 10 ng/mL PDGF-AA or PDGF-BB (R&D Systems) for the PDGFRα homodimer and PDGFRβ homodimer cell lines, respectively, and expanded to generate clonal cell lines. Final clones chosen for PDGFRα and PDGFRβ homodimer cell lines were confirmed by PCR amplification of the inserted sequences from genomic DNA using the following primers:

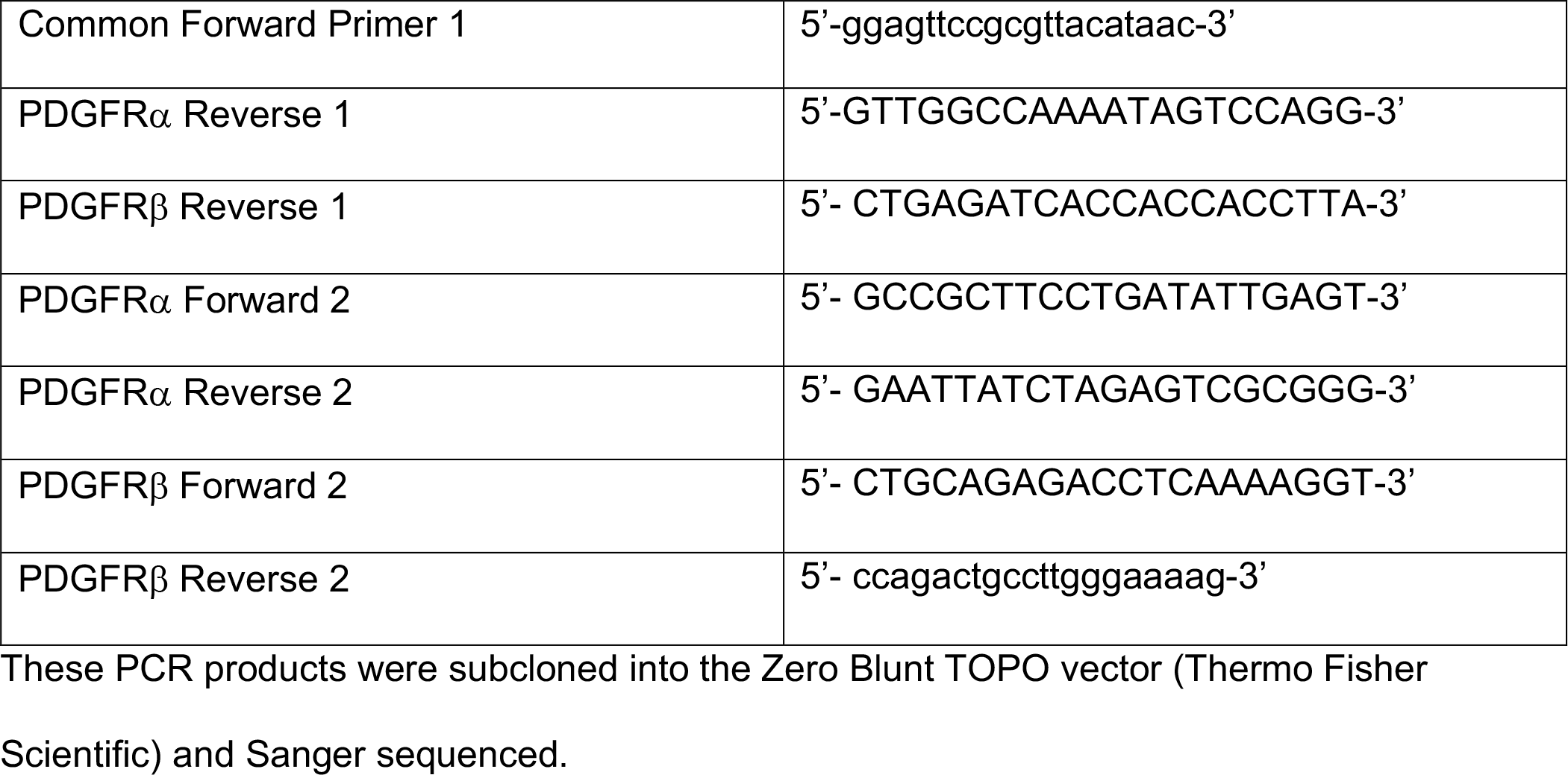

### Cell culture

PDGFR-BiFC stable cells were cultured in RPMI growth media [RPMI (Gibco) supplemented with 100 U/mL penicillin (Gibco), 100 µg/mL streptomycin (Gibco) containing 10% FBS (Hyclone Laboratories)] at 37°C in 5% carbon dioxide. Once the stable cell lines were established, they were split at a ratio of 1:5 for maintenance. PDGFRα homodimer cells were used for experiments at passage 6-15, and PDGFRβ homodimer cells were used for experiments at passage 10-18. When serum starved prior to ligand stimulation, cells were grown in HITES media [DMEM/F12 (Corning) supplemented with 0.1% bovine serum albumin (Fisher), 10 mM beta-estradiol (Sigma Aldrich), 10 mM hydrocortisone (Sigma Aldrich), 5 µg/mL insulin (Sigma Aldrich), 100 U/mL penicillin (Gibco), 100 µg/mL streptomycin (Gibco), NaHCO3 (Santa Cruz Biotechnology), 30 nM Na3SeO3 (Sigma Aldrich) and 10 µg/mL apo-transferrin (Sigma-Aldrich)].

### Immunoprecipitations and western blotting

PDGFR-BiFC-HCC15 cells were cultured as described above. To induce PDGFRα homodimer or PDGFRβ homodimer signaling, PDGFRα homodimer cells and PDGFRβ homodimer cells at ∼60-70% confluence were serum starved for 24 h in HITES media and stimulated with 10 ng/mL PDGF-AA or PDGF-BB ligand (R&D Systems), respectively, for the indicated length of time. To induce epidermal growth factor receptor (EGFR) signaling, cells were similarly stimulated with 10 ng/mL EGF (PeproTech, Inc.). When applicable, cells were pretreated with 10 µg/mL cycloheximide in DMSO (Sigma-Aldrich) or an equivalent volume of DMSO 30 min before PDGF ligand stimulation. Protein lysates for immunoprecipitation were generated by resuspending cells in ice-cold GFP-Trap lysis buffer [20 mM Tris-HCl (pH 7.5), 150 mM NaCl, 0.5% Nonidet P-40, 1 mM EDTA, 1× complete Mini protease inhibitor cocktail (Roche Diagnostics), 1 mM PMSF] and collecting cleared lysates by centrifugation at 13,400 g at 4°C for 20 min. For immunoprecipitations, cell lysates (500 µg) were incubated with GFP-Trap agarose beads (Bulldog-Bio) for 1 h at 4°C. Beads were washed three times with ice-cold GFP-Trap wash/dilution buffer [10 mM Tris HCl (pH 7.5), 150 mM NaCl, 0.5 M EDTA] and the precipitated proteins were eluted with Laemmli buffer containing 10% β-mercaptoethanol, heated for 10 min at 100°C and separated by SDS-PAGE. For western blotting analysis of whole cell lysates, protein lysates were generated by resuspending cells in ice-cold NP-40 lysis buffer [20 mM Tris-HCl (pH 8), 150 mM NaCl, 10% glycerol, 1% Nonidet P-40, 2 mM EDTA, 1× complete Mini protease inhibitor cocktail (Roche Diagnostics), 1 mM PMSF, 10 mM NaF, 1 mM Na3VO4, 25 mM β-glycerophosphate] and collecting cleared lysates by centrifugation at 13,400 g at 4°C for 20 min. Laemmli buffer containing 10% β-mercaptoethanol was added to the lysates, which were heated for 5 min at 100°C. Proteins were subsequently separated by SDS-PAGE. Western blot analysis was performed according to standard protocols using horseradish peroxidase-conjugated secondary antibodies. The following antibodies were used for western blotting: PDGFRα (1:200; C-20; sc338; Santa Cruz Biotechnology) (validation, dimerization and receptor degradation experiments); PDGFRβ (1:200; 958; sc432; Santa Cruz Biotechnology) (validation, dimerization and receptor degradation experiments); β-tubulin (1:1000; E7; E7; Developmental Studies Hybridoma Bank); phospho-PDGFRα (Tyr 849)/PDGFRβ (Tyr857) (1:1000; C43E9; 3170; Cell Signaling Technology); PDGFRα (1:1000; D13C6; 5241; Cell Signaling Technology) (autophosphorylation experiments); PDGFRβ (1:1000; 28E1; 3169; Cell Signaling Technology) (autophosphorylation experiments); phospho-Erk1/2 (1:1000; Thr202/Thr204; 9101; Cell Signaling Technology); Erk1/2 (1:1000; 9102; Cell Signaling Technology); phospho-Akt (1:1000; Ser473; 9271; Cell Signaling Technology); Akt (1:1000; 9272; Cell Signaling Technology); horseradish peroxidase-conjugated goat anti-rabbit IgG (1:20,000; 111035003; Jackson ImmunoResearch Laboratories); horseradish peroxidase-conjugated goat anti-mouse IgG (1:20,000; 115035003; Jackson ImmunoResearch laboratories). Quantifications of signal intensity were performed with ImageJ software (version 1.52a, National Institutes of Health). Relative dimerized PDGFR levels were determined by normalizing GFP-Trap immunoprecipitated PDGFR levels to total PDGFR levels. Relative phospho-PDGFR levels were determined by normalizing to total PDGFR levels. Relative degraded PDGFR levels were determined by normalizing cycloheximide-treated PDGFR levels to DMSO-treated PDGFR levels. Relative phospho-ERK1/2 levels were determined by normalizing to total ERK1/2 levels. Relative phospho-AKT levels were determined by normalizing to total AKT levels. When applicable, statistical analyses were performed with Prism 9 (GraphPad Software) using a ratio paired *t*-test within each cell line (comparing individual ligand treatment timepoint values to the no ligand 0 min timepoint value) and a two-tailed, unpaired *t*-test with Welch’s correction between each cell line. Immunoprecipitation and western blotting experiments were performed across at least three independent experiments.

### Immunofluorescence analysis

Cells were seeded onto glass coverslips coated with 5 µg/mL human plasma fibronectin purified protein (EMD Millipore Corporation) at a density of 80,000 cells and 40,000 cells for the PDGFRα homodimer cell line and PDGFRβ homodimer cell line, respectively, in RPMI growth media. 24 h later, cells were washed with 1x phosphase buffered saline (PBS) and serum starved in HITES media. HITES media was replaced 23 h later. After 54 min, coverslips were photobleached for 1 min with an Axio Observer 7 fluorescence microscope (Carl Zeiss) using the 2.5x objective and 488 nm laser. Cells were allowed to recover for 5 min and were treated with 10 ng/mL PDGF-AA or PDGF-BB ligand (R&D Systems) for the indicated amount of time. Cells were fixed in 4% paraformaldehyde in PBS with 0.1% Triton X-100 for 10 min and washed in PBS. Cells were blocked for 1 h in 5% normal donkey serum (Jackson ImmunoResearch Laboratories) in PBS and incubated overnight at 4°C in primary antibody diluted in 1% normal donkey serum in PBS. After washing in PBS, cells were incubated in Alexa Fluor 546-conjugated donkey anti-rabbit secondary antibody (1:1,000; A21206; Invitrogen) or Alexa Fluor 546-conjugated donkey anti-mouse secondary antibody (1:1,000; A10036; Invitrogen) diluted in 1% normal donkey serum in PBS with 2 µg/mL 4’,6-diamidino-2-phenylindole (DAPI; Sigma-Aldrich) for 1 h. Cells were mounted in either VECTASHIELD HardSet Antifade Mounting Medium or VECTASHIELD Vibrance Antifade Mounting Medium (Vector Laboratories) and photographed using an Axiocam 506 mono digital camera (Carl Zeiss) fitted onto an Axio Observer 7 fluorescence microscope (Carl Zeiss) with the 63x oil objective with a numerical aperature of 1.4 at room temperature. The following antibodies were used for immunofluorescence analysis: PDGFRα (1:100; RB-1691; Thermo Fisher Scientific); PDGFRβ (1:25; 28D4; 558820; Becton Dickinson); Na^+^/K^+^-ATPase (1:500; EP1845Y; ab76020; Abcam); Rab5 (1:200; C8B1; 3547; Cell Signaling); Rab7 (1:100; D95F2; 9367; Cell Signaling); Rab4 (1:200; ab13252; Abcam); Rab11 (1:100; D4F5; 5589; Cell Signaling).

For fluorescence intensity measurements, at least 40 images with Z-stacks (0.24 µm between Z-stacks with a range of 3-15 Z-stacks) were taken per cell line across two trials per experiment. For marker co-localization experiments, at least 20 images with Z-stacks (0.24 µm between Z-stacks with a range of 1-15 Z-stacks) were taken per cell line per timepoint across two trials per experiment. Images were deconvoluted using ZEN Blue software (Carl Zeiss) using the “Better, fast (Regularized Inverse Filter)” setting. For all images, extended depth of focus was applied to Z-stacks using ZEN Blue software (Carl Zeiss) to generate images with the maximum depth of field. For fluorescence intensity measurements, background was subtracted using rolling background subtraction with a radius of 30 pixels using Fiji software (version 2.1.0/1.53c; National Institutes of Health). A region of interest (ROI) was drawn around each Venus-positive cell, and integrated density was measured and recorded as the fluorescence intensity. For marker co-localization measurements, an ROI was drawn around each Venus-positive cell in the corresponding Cy3 (marker) channel using Fiji software (version 2.1.0/1.53c; National Institutes of Health). For each image with a given ROI, the Cy3 channel and the EGFP channel were converted to 8-bit images. Co-localization was measured using the Colocalization Threshold function, where the rcoloc value (Pearson’s correlation coefficient (PCC)) was used in statistical analysis. Statistical analyses were performed on individual cell values with Prism 9 (GraphPad Software) using a two-tailed, unpaired *t*-test with Welch’s correction.

### Anchorage-independent growth assays

For measurement of anchorage-independent cell growth, 25,000 cells were suspended in 1.5 mL RPMI containing 100 U/mL penicillin (Gibco), 100 µg/mL streptomycin (Gibco), 10% FBS and 0.35% Difco Agar Noble (Becton Dickinson) and overlaid on a base layer containing 1.5 mL RPMI containing 100 U/mL penicillin (Gibco), 100 µg/mL streptomycin (Gibco), 10% FBS and 0.5% agar noble in 6-well plates. A feeding layer of 2 mL RPMI growth media supplemented with 10 ng/mL PDGF-AA or PDGF-BB ligand (R&D Systems) was added on top of the agar and replaced every day. Plates were incubated at 37°C in 5% carbon dioxide for 10 days, and viable colonies were stained overnight with 1 mg/mL Nitrotetrazoleum Blue (Sigma-Aldrich) in PBS at 37°C. Following a second overnight incubation at 4°C, wells were photographed using a COOLPIX S600 digital camera (Nikon). Images were made binary using Adobe Photoshop (version 21.0.3; Adobe), and colony number and area were quantified using Meta-Morph imaging software (Molecular Devices). Statistical analyses were performed on values from individual images with Prism 9 (GraphPad Software) using a two-tailed, unpaired *t*-test with Welch’s correction.

### Transwell assays

Cells were serum-starved for 24 h in HITES media. Cell culture inserts for 24-well plates containing polyethylene terephthalate membranes with 8 µm pores (Corning Inc.) were coated with 5 µg/mL human plasma fibronectin purified protein (EMD Millipore Corporation). Cells were seeded at a density of 315,000 cells per insert in 250 µL HITES media, and inserts were immersed in 500 µL RPMI media containing 10% FBS in the absence or presence of 10 ng/mL PDGF-AA or PDGF-BB ligand (R&D Systems) for 24 h. Migrated cells were subsequently fixed in 4% PFA in PBS for 10 min and stained in 0.1% crystal violet in 10% ethanol for 10 min.

Dried inserts were photographed using an Axiocam 105 color camera fitted onto a Stemi 508 stereo microscope with the 0.5x objective at room temperature (Carl Zeiss). Five fields of cells from each of six independent trials were photographed and quantified by measuring integrated density with ImageJ software (version 1.52a; National Institutes of Health). Statistical analyses were performed on values from individual images with Prism 9 (GraphPad Software) using a two-tailed, unpaired *t*-test with Welch’s correction.

### Online supplemental material

**Fig. S1** shows a schematic describing the PDGFR-BiFC approach, images of DAPI-stained nontransduced HCC15 cells in the absence or presence of PDGF ligand stimulation and western blots of whole cell lysates depicting appropriate PDGFR expression in the two PDGFR homodimer cell lines. **Fig. S2** shows a bar graph depicting PCC of Venus signal and Na^+^/K^+^-ATPase at 90 min of ligand treatment and corresponding immunofluorescence data. **Fig. S3** shows phospho-ERK1/2 and phospho-AKT western blots of whole cell lysates of both PDGFR homodimer cell lines in the absence or presence of EGF ligand stimulation with corresponding quantifications.

## Results

### Generation and validation of PDGFR-BiFC stable cell lines

To investigate PDGFR dimer-specific dynamics, we generated stable cell lines expressing BiFC-tagged PDGFRs. We cloned plasmids expressing C-terminal protein fusions of each PDGFR with BiFC fragments corresponding to the N-terminal (V1) or C-terminal (V2) regions of the Venus fluorescent protein (**Fig. S1A**). Using lentiviral transduction of HCC15 cells (**Fig. S1, B-C’**), we stably integrated PDGFR-BiFC sequences (PDGFRα-V1/PDGFRα-V2 and PDGFRβ-V1/PDGFRβ-V2) to generate two cell lines: a PDGFRα homodimer cell line and a PDGFRβ homodimer cell line, respectively. We confirmed that the relevant sequences were inserted into the genome and established that the relevant PDGFR was expressed in each cell line (**Fig. S1, D and E**). HCC15 cells were selected based on the lack of PDGFR expression, as assessed by RNA-sequencing, as the presence of endogenous PDGFRs would confound the interpretation of BiFC events. Further, these cells have little to no PDGF ligand expression, such that the BiFC-tagged PDGFRs would only be activated upon exogenous ligand stimulation. Finally, HCC15 cells express all of the relevant adaptor proteins and signaling molecules known to function downstream of PDGFR activation (Barretina et al., 2012; Cancer Cell Line Encyclopedia and Genomics of Drug Sensitivity in Cancer, 2015; Heldin and Westermark, 1999). Within these stable cell lines, the individual N- and C-terminal fragments of the Venus fluorescent protein are expected to be non-fluorescent when bound to monomeric receptors. However, upon receptor dimerization, the N- and C-terminal fragments should co-localize, resulting in a functional Venus fluorescent protein. This BiFC event can be visualized through fluorescence microscopy or purified biochemically utilizing a GFP-Trap nanobody, which has an epitope that spans V1 and V2 (Croucher et al., 2016).

We confirmed the BiFC event upon exogenous ligand treatment of both cell lines through fluorescence microscopy (**Fig. 1, A-D’**). We stimulated PDGFRα homodimers with PDGF-AA ligand and PDGFRβ homodimers with PDGF-BB ligand, as previous data indicated that PDGFRα homodimers are primarily responsive to activation by PDGF-AA, while PDGFRβ homodimers are primarily activated by PDGF-BB (Bostrom et al., 1996; Fantauzzo and Soriano, 2016; Leveen et al., 1994; Lokker et al., 1997; Soriano, 1994; Soriano, 1997). After 24 h of starvation in HITES media, which lacks any growth factors, we photobleached the coverslips to ensure that all imaging captured newly-formed BiFC events. We then stimulated with PDGF ligand for 5 min and observed Venus fluorescence in both the PDGFRα homodimer and the PDGFRβ homodimer cell lines (**Fig. 1, B, B’, D, D’**). At this timepoint, Venus expression in the PDGFRα homodimer cell line was typically observed as small, internalized puncta, whereas in the PDGFRβ homodimer cell line, Venus expression was at both the cell membrane and inside the cell. Quantification of fluorescence intensity revealed a significant increase in Venus intensity upon 5 min of ligand treatment for the two cell lines (PDGFRαV1/V2: *p* < 0.0001, PDGFRβV1/V2: *p* = 0.0002; **Fig. 1, E and F**). Importantly, fluorescence intensity was comparable between the two cell lines (PDGFRαV1/V2: 21.19 ± 1.958 arbitrary units (A.U.); PDGFRβV1/V2: 19.32 ± 1.564 A.U.; **Fig. 1, E and F**).

**Figure 1.**
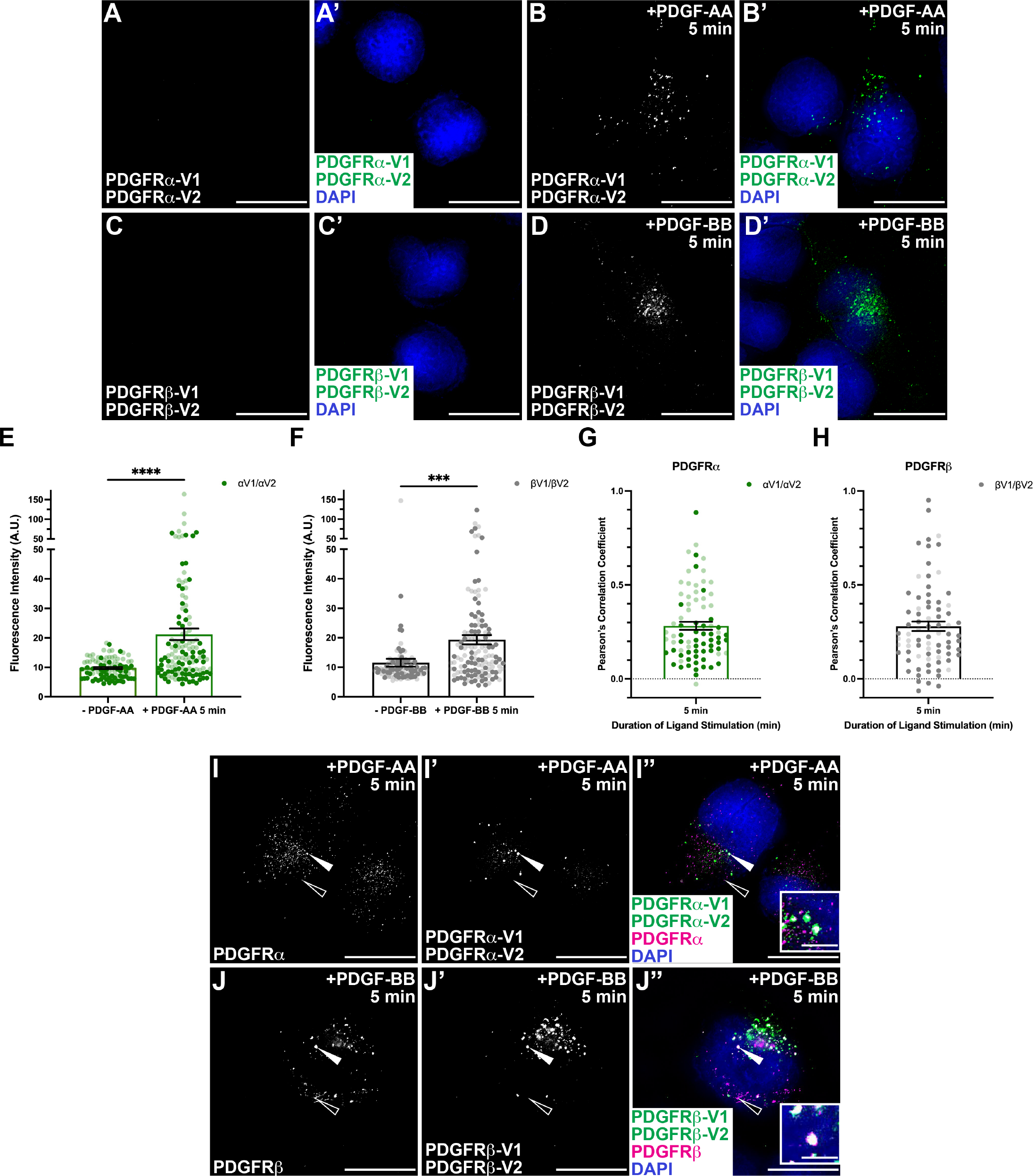
Validation of PDGFR-BiFC stable cell lines. (A-D’) Venus expression (white/green) as assessed by fluorescence analysis of HCC15 cells transduced with PDGFRα-V1 and PDGFRα-V2 (A-B’) or PDGFRβ-V1 and PDGFRβ-V2 (C-D’) in the absence (A, A’, C, C’) or presence (B, B’, D, D’) of PDGF ligand for 5 min. Nuclei were stained with DAPI (blue; A’-D’). (E-F) Bar graphs depicting fluorescence intensity for PDGFRα homodimer (E) and PDGFRβ homodimer (F) cell lines. Data are mean ± SEM. Statistical analyses were performed using a two-tailed, unpaired *t*-test with Welch’s correction. ***, p<0.001; ****, p<0.0001. Colored circles correspond to independent experiments. *n*>40 technical replicates across each of 2 biological replicates. (G-H) Bar graphs depicting Pearson’s correlation coefficient of PDGFRα homodimer cell line with an anti-PDGFRα antibody (G) and PDGFRβ homodimer cell line with an anti-PDGFRβ antibody (H) following PDGF ligand stimulation for 5 min. Data are mean ± SEM. Colored circles correspond to independent experiments. *n*>20 technical replicates across each of 2 biological replicates. (I-J”) PDGFR antibody expression (white/magenta; I, I”, J, J”) and/or Venus expression (white/green; I’, I”, J’, J”) as assessed by (immuno)fluorescence analysis of PDGFRα homodimer (I-I”) and PDGFRβ homodimer (J-J”) cell lines. Insets in I” and J” are regions where white arrows are pointing. Nuclei were stained with DAPI (blue; I”, J”). White arrows denote co-localization; white outlined arrows denote lack of co-localization. Scale bars, 20 µm. Inset scale bars, 3 µm.

Next, we investigated co-localization of the Venus signal with signal from anti-PDGFR antibodies. These data revealed that Venus signal co-localized with a subset of PDGFR expression at 5 min of ligand stimulation (**Fig. 1, G-J”**). Pearson’s correlation coefficient (PCC) for the Venus signal and PDGFR expression was similar between the PDGFRα homodimer (0.2822 ± 0.0211 PCC) and the PDGFRβ homodimer cell lines (0.2800 ± 0.0253 PCC; **Fig. 1, G and H**). The regions of receptor expression that were not Venus-positive likely represent monomeric receptors and/or potential V1/V1 or V2/V2 dimers, which would not be expected to fluoresce (**Fig. 1, I-J”**). Importantly, the fact that we observed less than complete positive co-localization (PCC values <1) indicated that the presence of the BiFC fragments alone did not drive dimerization.

### PDGFRβ receptors homodimerize more quickly than PDGFRα receptors

We further confirmed the BiFC event upon ligand treatment for both cell lines biochemically. Cells for both PDGFR homodimers were stimulated with PDGF ligand for 2, 5 and 15 min. Subsequent immunoprecipitation with the GFP-Trap nanobody was followed by western blotting for each receptor, normalizing the immunoprecipitation signal to the whole cell lysate signal. This revealed an increase in dimerized receptors following ligand treatment of both cell lines, with peak dimerization occurring at 5 min for both PDGFRα homodimers (1.245 ± 0.0361 relative induction (R.I.)) and PDGFRβ homodimers (1.492 ± 0.3608 R.I.; **Fig. 2, A-C**). Furthermore, PDGFRβ homodimers appeared to dimerize more quickly than PDGFRα homodimers, reaching near peak dimerization at 2 min of ligand treatment (1.473 ± 0.2682 R.I.; **Fig. 2C**).

**Figure 2.**
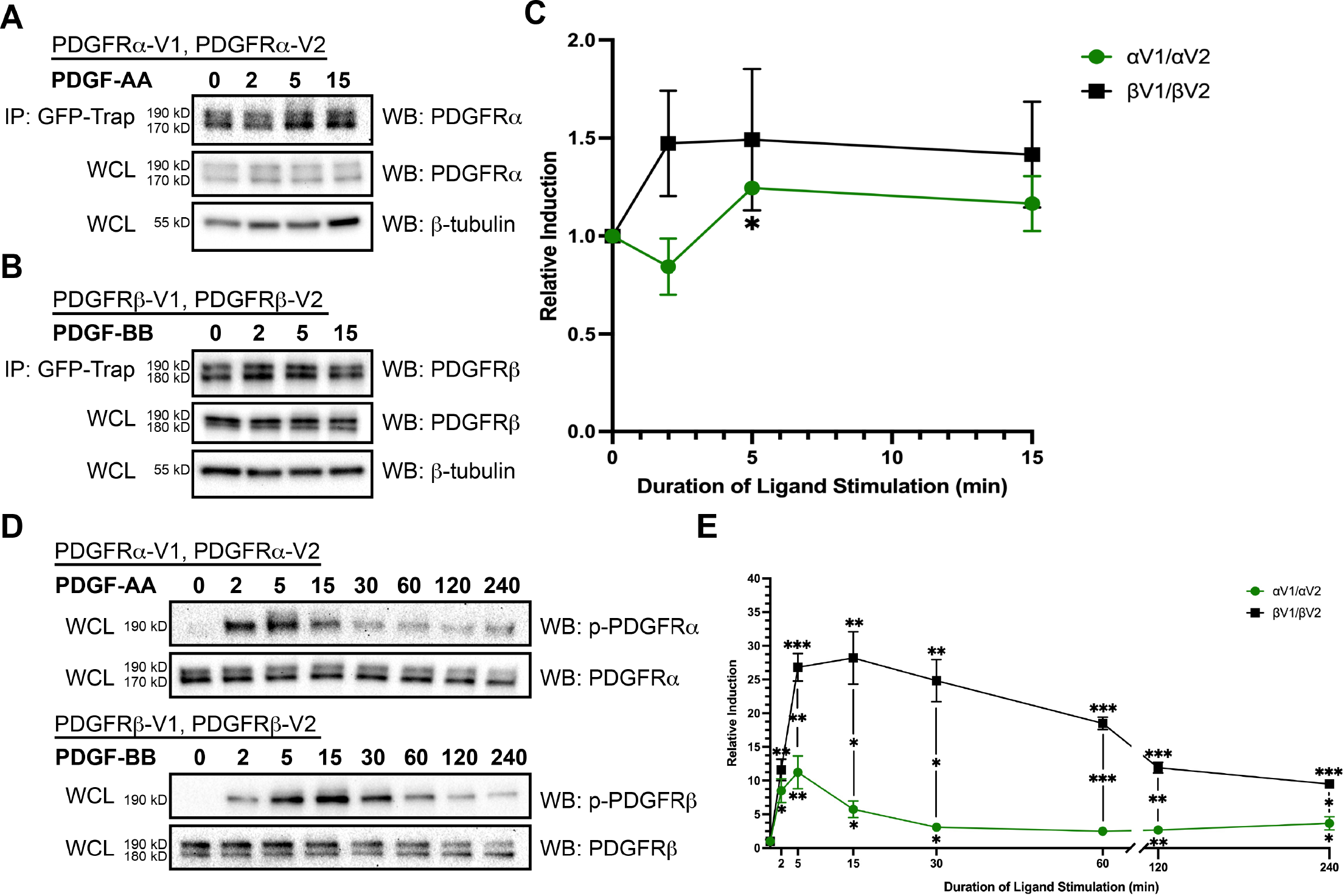
PDGFRβ receptors homodimerize more quickly than alpha receptors with increased levels of auto-phosphorylation. (A, B) Immunoprecipitation (IP) of dimerized PDGFRα receptors (A) and PDGFRβ receptors (B) with GFP-Trap nanobody from cells that were treated with PDGF ligand for 2-15 min followed by western blotting (WB) with anti-PDGFRα (A) or PDGFRβ (B) antibodies. (C) Line graph depicting quantification of band intensities from *n*=3 biological replicates as in A and B. Data are mean ± SEM. Statistical analyses were performed using a ratio paired *t*-test within each cell line and a two-tailed, unpaired *t*-test with Welch’s correction between each cell line. *, p<0.05. (D) Western blot analysis of whole cell lysates (WCL) from PDGFRα homodimer (top) and PDGFRβ homodimer (bottom) cell lines following a timecourse of PDGF stimulation from 2 min to 4 h with an anti-phospho-PDGFR antibody. (E) Line graph depicting quantification of band intensities from *n*=3 biological replicates as in D. Data are mean ± SEM. Statistical analyses were performed using a ratio paired *t*-test within each cell line and a two-tailed, unpaired *t*-test with Welch’s correction between each cell line. *, p<0.05; **, p<0.01; ***, p<0.001.

Next, we investigated the activation of both PDGFR homodimers by assessing receptor autophosphorylation following a timecourse of ligand stimulation. Each cell line was stimulated with PDGF ligand from 2 min to 4 h. Western blotting of whole cell lysates using an anti-phospho-PDGFR antibody revealed that both PDGFR homodimers were significantly and robustly autophosphorylated upon ligand treatment. Peak activation occurred at 5 min of ligand stimulation for PDGFRα homodimers (11.24 ± 2.437 R.I.) and at 15 min for PDGFRβ homodimers (28.19 ± 3.895 R.I.; **Fig. 2, D and E**). Critically, there was very little to no activation of the PDGFRs in the absence of ligand (**Fig. 2D**), consistent with previous findings that both ligand binding and dimerization are required for receptor activation (Kelly et al., 1991; Yang et al., 2010). Furthermore, PDGFRβ homodimers exhibited more robust autophosphorylation than PDGFRα homodimers from 5 min to 4 h of PDGF ligand stimulation (**Fig. 2, D and E**). Together, these results suggested that PDGFRβ receptors homodimerize more quickly and have increased levels of autophosphorylation than PDGFRα homodimers.

### PDGFRα homodimers are trafficked more quickly and degraded faster, while PDGFRβ homodimers are more likely to be recycled

It is well-established that PDGFRs dimerize at the cell membrane upon ligand binding (Herren et al., 1993; Shim et al., 2010) and further, that PDGFR dimers become internalized and trafficked in the cell following activation (De Donatis et al., 2008; Disanza et al., 2009; Miaczynska, 2013; Pahara et al., 2010; Rogers and Fantauzzo, 2020). As mentioned above, the Venus signal in the two cell lines exhibited different subcellular localization following PDGF ligand treatment. To further investigate this, we evaluated membrane localization of the PDGFR dimers at early timepoints of PDGF ligand treatment by measuring co-localization of the Venus signal with signal from an antibody recognizing the membrane marker Na^+^/K^+^-ATPase. We observed relatively similar levels of co-localization for PDGFRα homodimers and PDGFRβ homodimers at 1 and 2 min following PDGF ligand stimulation (**Fig. 3A**). At 5 min of ligand treatment, however, PDGFRβ homodimers displayed significantly greater co-localization (0.2283 ± 0.0262 PCC) with Na^+^/K^+^-ATPase than PDGFRα homodimers (0.1641 ± 0.0186 PCC; *p* = 0.0478; **Fig. 3, A-C”**). To assess whether PDGFR dimer internalization could account for this comparative decrease in membrane localization for PDGFRα homodimers, we next measured the co-localization of the Venus signal with signal from an antibody recognizing the early endosome marker Rab5 (Zerial and McBride, 2001). These analyses revealed that PDGFRα homodimers co-localized significantly more with Rab5 at 2 (0.2598 ± 0.0261 PCC), 5 (0.1562 ± 0.0122 PCC) and 10 min (0.1630 ± 0.0192 PCC) of PDGF ligand treatment than PDGFRβ homodimers (0.1102 ± 0.0165 PCC; *p* < 0.0001; 0.0951 ± 0.0189 PCC; *p* = 0.0076; 0.0781 ± 0.0174 PCC; *p* = 0.0013, respectively), indicating a greater likelihood for PDGFRα homodimers to be trafficked at these timepoints (**Fig. 3, D-F”**). Collectively, these data suggested that PDGFRα homodimers are trafficked more quickly, while PDGFRβ homodimers dwell at the membrane longer.

**Figure 3.**
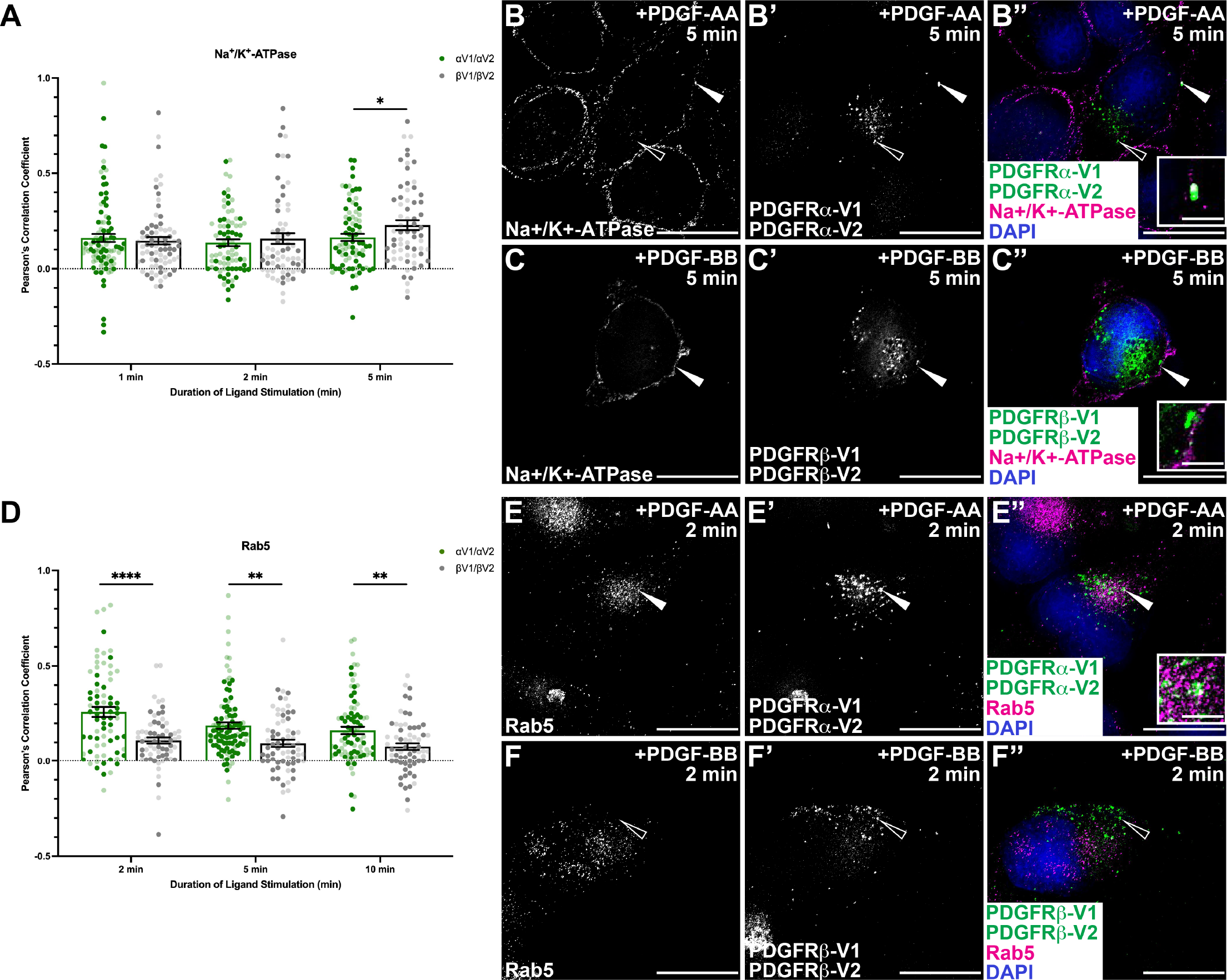
PDGFRα homodimers are trafficked more quickly than PDGFRβ homodimers. (A, D) Bar graphs depicting Pearson’s correlation coefficient of the PDGFRα homodimer and PDGFRβ homodimer cell lines with an anti-Na^+^/K^+^-ATPase antibody (A) or an anti-Rab5 antibody (D) following PDGF ligand stimulation from 1-5 min (A) or 2-10 min (D). Data are mean ± SEM. Statistical analyses were performed using a two-tailed unpaired *t*-test with Welch’s correction. *, p<0.05; **, p<0.01; ****, p<0.0001. Colored circles correspond to independent experiments. *n*>20 technical replicates across each of 2 biological replicates. (B-C”, E-F”) Na^+^/K^+^-ATPase antibody expression (white/magenta; B, B”, C, C”) or Rab5 antibody expression (white/magenta; E, E”, F, F”) and/or Venus expression (white/green; B’, B”, C’, C”, E’, E”, F’, F”) as assessed by (immuno)fluorescence analysis of PDGFRα homodimer (B-B”, E-E”) and PDGFRβ homodimer (C-C”, F-F”) cell lines. Insets in B”, C” and F” are regions where white arrows are pointing. Nuclei were stained with DAPI (blue; B”, C”, E”, F”). White arrows denote co-localization; white outlined arrows denote lack of co-localization. Scale bars, 20 µm. Inset scale bars, 3 µm.

We next assessed trafficking of the PDGFR homodimers through late endosomes, which precedes trafficking to the lysosome for degradation (Mellman, 1996), by examining co-localization of the Venus signal with signal from an antibody recognizing the late endosome marker Rab7 (Zerial and McBride, 2001). These data revealed that PDGFRα homodimers co-localized significantly more with Rab7 (0.3271 ± 0.0165 PCC) than PDGFRβ homodimers (0.2592 ± 0.0211 PCC; *p* = 0.0125) at 90 min of ligand treatment (**Fig. 4, A-C”**). This finding suggested that PDGFRα homodimers may be degraded more quickly and/or to a greater extent than PDGFRβ homodimers. To investigate this late endosome trafficking further, we biochemically evaluated receptor degradation following PDGF ligand stimulation. We performed a 30 min pretreatment with cycloheximide to inhibit translation of new receptors during the timecourse. Cells were then stimulated with PDGF ligand from 2 min to 4 h. We collected cell lysates and performed western blotting for PDGFR levels in each condition. These data revealed a sharp decrease in overall receptor levels for both PDGFR homodimers. Specifically, following 4 h of ligand treatment, we observed 80% and 78% decreases from initial receptor levels for PDGFRα homodimers (*p* = 0.0316) and PDGFRβ homodimers, respectively, indicating a similar extent of degradation by the end of the timecourse (**Fig. 4, D and E**). Importantly, we observed that PDGFRα homodimers were degraded at a quicker rate, particularly between the 60 and 120 min timepoints (**Fig. 4E**). Taken together, these data suggested that PDGFRα homodimers are degraded more quickly than PDGFRβ homodimers.

**Figure 4.**
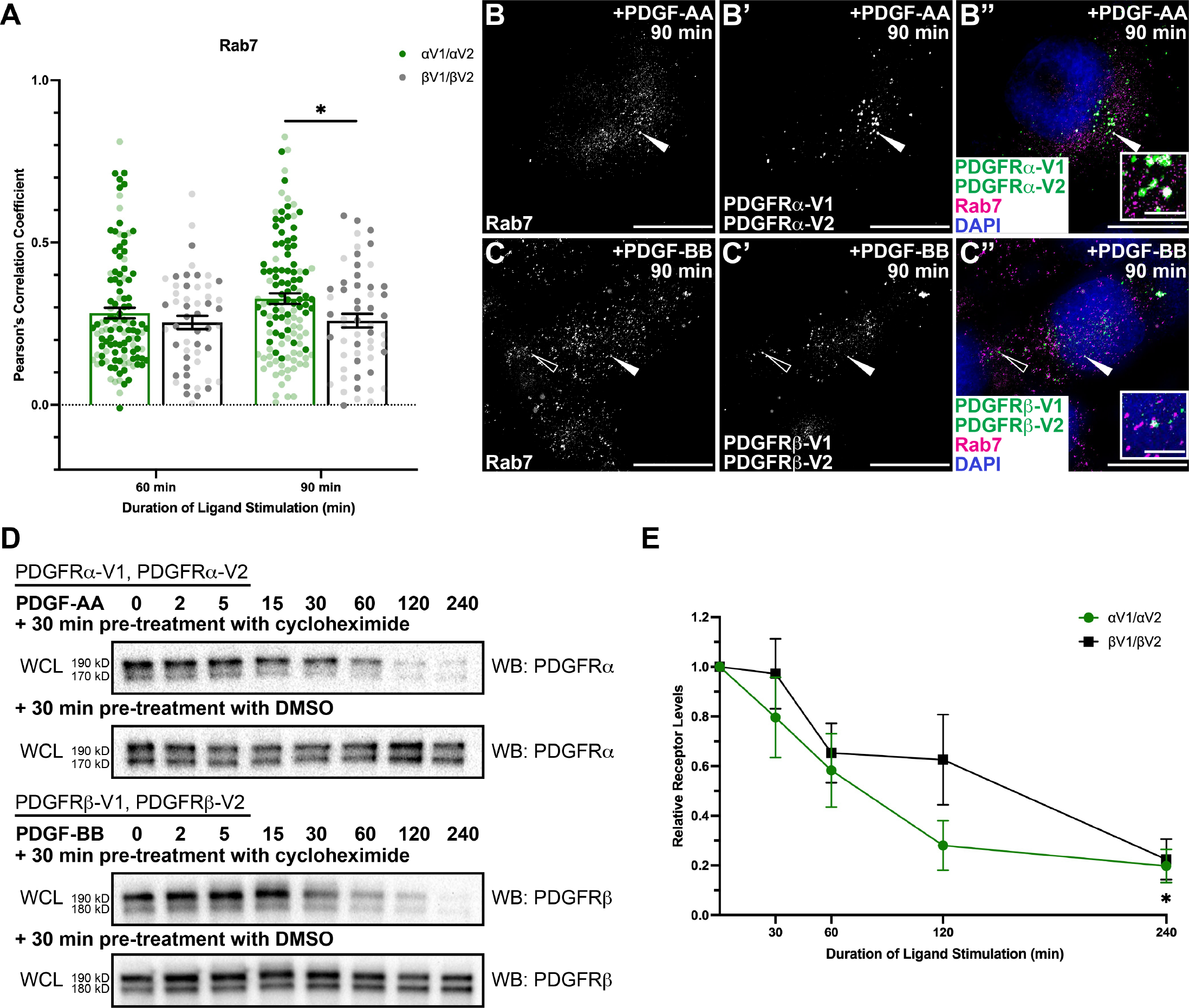
PDGFRα homodimers are degraded more quickly than PDGFRβ homodimers. (A) Bar graph depicting Pearson’s correlation coefficient of the PDGFRα homodimer and PDGFRβ homodimer cell lines with an anti-Rab7 antibody following PDGF ligand stimulation from 60-90 min. Data are mean ± SEM. Statistical analyses were performed using a two-tailed unpaired *t*-test with Welch’s correction. *, p<0.05. Colored circles correspond to independent experiments. *n*>20 technical replicates across each of 2 biological replicates. (B-C”) Rab7 antibody expression (white/magenta; B, B”, C, C”) and/or Venus expression (white/green; B’, B”, C’, C”) as assessed by (immuno)fluorescence analysis of PDGFRα homodimer (B-B”) and PDGFRβ homodimer (C-C”) cell lines. Insets in B” and C” are regions where white arrows are pointing. Nuclei were stained with DAPI (blue; B”, C”). White arrows denote co-localization; white outlined arrows denote lack of co-localization. Scale bars, 20 µM. Inset scale bars, 3 µm. (D) Western blot (WB) analysis of whole cell lysates (WCL) from PDGFRα homodimer (top) and PDGFRβ homodimer (bottom) cell lines following pre-treatment with 10 µg/mL cycloheximide or equivalent volume of DMSO and a timecourse of PDGF stimulation from 2 min to 4 h with anti-PDGFRα (top) or anti-PDGFRβ (bottom) antibodies. (E) Line graph depicting quantification of band intensities from *n*=3 biological replicates as in D. Data are mean ± SEM. Statistical analyses were performed using a ratio paired *t*-test within each cell line and a two-tailed, unpaired *t*-test with Welch’s correction between each cell line. *, p<0.05.

In contrast to being degraded, PDGFRs can alternatively be recycled to the membrane for continued signaling. To examine receptor recycling, we first quantified the co-localization of the Venus signal with signal from an antibody recognizing the rapid recycling endosome marker Rab4 (Zerial and McBride, 2001). Quantification indicated that PDGFRβ homodimers co-localized significantly more with Rab4 (0.2353 ± 0.0262 PCC) than PDGFRα homodimers (0.1593 ± 0.0159 PCC; *p* = 0.0145) at 15 min of ligand treatment (**Fig. 5, A-C”**). Next, we examined co-localization of the Venus signal with signal from an antibody recognizing the slow recycling endosome marker Rab11 (Zerial and McBride, 2001). This analysis revealed that PDGFRβ homodimers co-localized significantly more with Rab11 (0.1829 ± 0.0228 PCC, 0.1925 ± 0.0223 PCC, respectively) than PDGFRα homodimers (0.1131 ± 0.0113 PCC; *p* = 0.0075; 0.1189 ± 0.0148; *p =* 0.0067, respectively) at both 60 and 90 min of PDGF treatment (**Fig. 5, D-F”**). Consistently, at 90 min of PDGF ligand stimulation, PDGFRβ homodimers co-localized significantly more with the membrane marker Na^+^/K^+^-ATPase (0.1433 ± 0.0186 PCC) than PDGFRα homodimers (0.0965 ± 0.0133 PCC; *p* = 0.0429; **Fig. S2, A-C”**). Collectively, these data demonstrated that PDGFRβ homodimers are more likely to be recycled back to the cell membrane through these rapid and slow recycling routes than PDGFRα homodimers.

**Figure 5.**
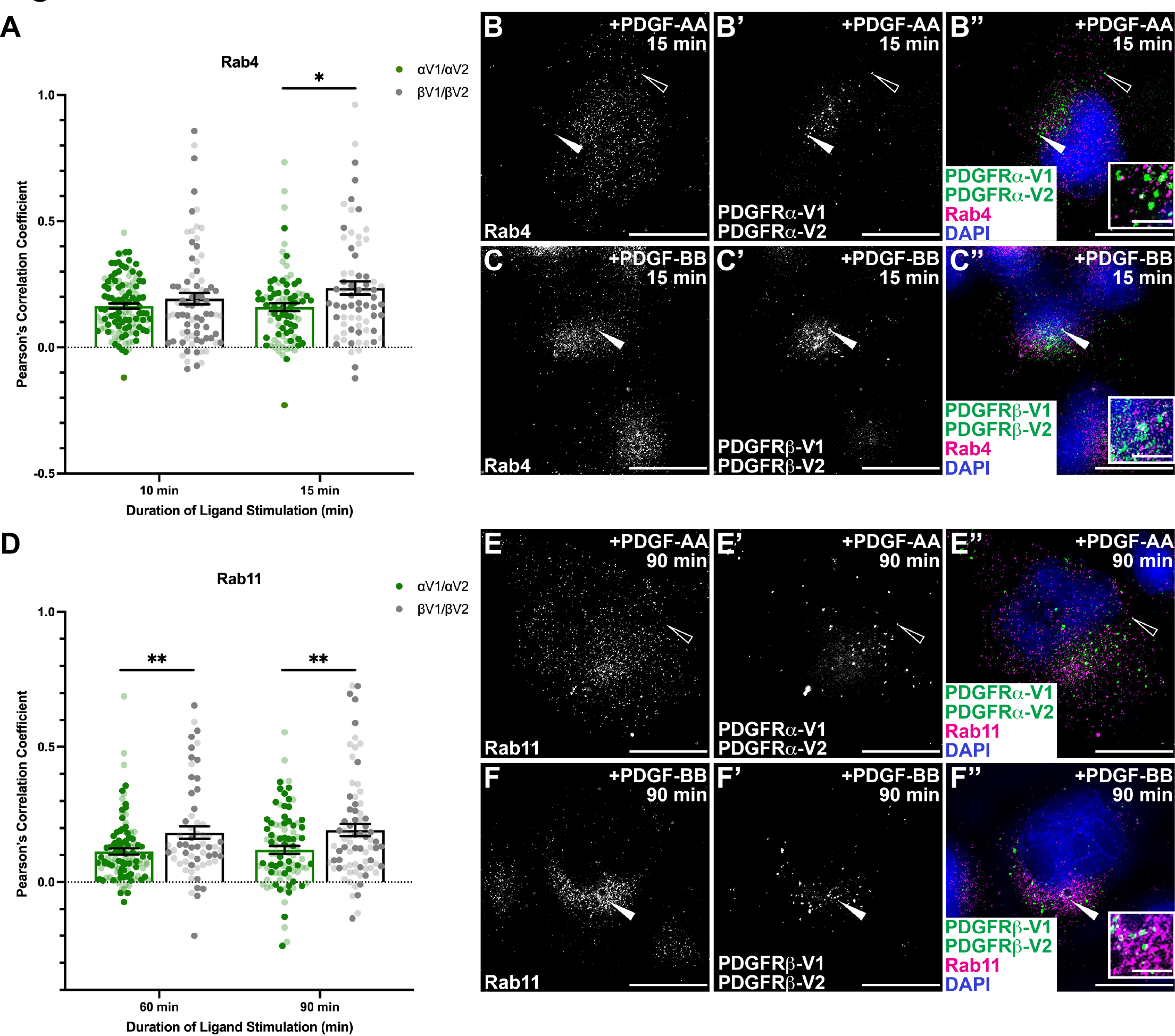
PDGFRβ homodimers are more likely to be recycled back to the cell membrane than PDGFRα homodimers. (A, D) Bar graph depicting Pearson’s correlation coefficient of the PDGFRα homodimer and PDGFRβ homodimer cell lines with an anti-Rab4 antibody (A) or an anti-Rab11 antibody (D) following PDGF ligand stimulation from 10-15 min (A) or 60-90 min (D). Data are mean ± SEM. Statistical analyses were performed using a two-tailed unpaired *t*-test with Welch’s correction. *, p<0.05; **, p<0.01. Colored circles correspond to independent experiments. *n*>20 technical replicates across each of 2 biological replicates. (B-C”, E-F”) Rab4 antibody expression (white/magenta; B, B”, C, C”) or Rab11 antibody expression (white/magenta; E, E”, F, F”) and/or Venus expression (white/green; B’, B”, C’, C”, E’, E”, F’, F”) as assessed by (immuno)fluorescence analysis of PDGFRα homodimer (B-B”, E-E”) and PDGFRβ homodimer (C-C”, F-F”) cell lines. Insets in B”, C” and F” are regions where white arrows are pointing. Nuclei were stained with DAPI (blue; B”, C”, E”, F”). White arrows denote co-localization; white outlined arrows denote lack of co-localization. Scale bars, 20 µm. Inset scale bars, 3 µm.

### PDGFRβ homodimer activation induces a greater amplitude of downstream signaling

We postulated that the differences in dimerization, activation and trafficking between PDGFR homodimers may affect downstream cell signaling. We performed a timecourse of ligand stimulation for each PDGFR homodimer cell line from 2 min to 4 h followed by western blotting for two well-studied effector molecules downstream of PDGFR activation: ERK1/2 and AKT (Fantauzzo and Soriano, 2016; Lemmon and Schlessinger, 2010; Vasudevan et al., 2015). Phosphorylation of ERK1/2 in response to PDGFRα homodimer signaling reached an initial peak at 5 min following PDGF ligand stimulation (1.469 ± 0.4787 R.I.) and remained relatively stable throughout the timecourse (1.325 ± 0.0693 R.I.; **Fig. 6, A and B**). Phosphorylation of ERK1/2 in response to PDGFRβ homodimer signaling displayed a similar curve to that of PDGFRα homodimers but peaked at 30 min of PDGF stimulation (2.318 ± 0.5807 R.I.) with a much higher amplitude than the PDGFRα homodimers (**Fig. 6, A and B**). Phosphorylation of AKT in response to PDGFRα homodimer activation initially peaked at 15 min of ligand treatment (2.038 ± 0.2840 R.I.; *p* = 0.0382) and, comparable to the phosphorylation dynamics of ERK1/2, remained relatively stable (**Fig. 6, C and D**).

**Figure 6.**
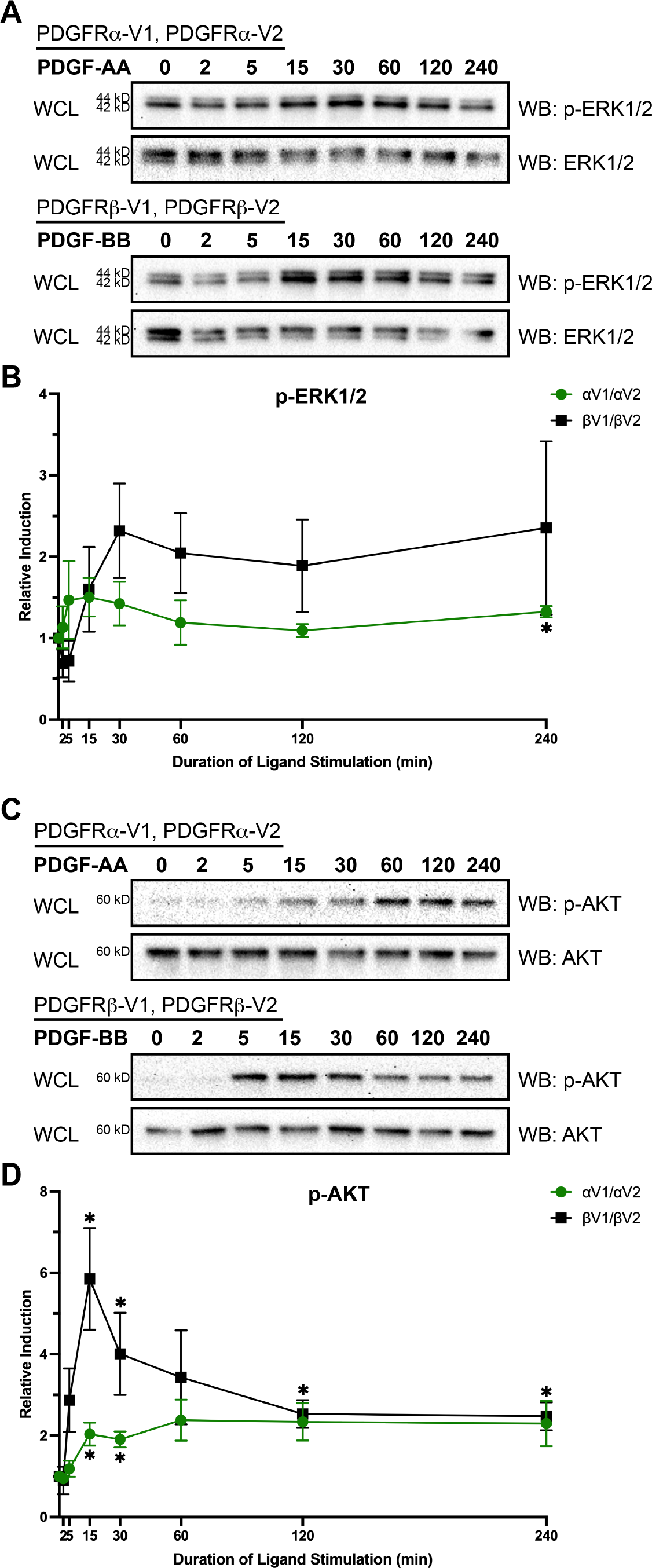
PDGFRβ homodimer activation induces a greater amplitude of downstream signaling. (A, C) Western blot (WB) analysis of whole cell lysates (WCL) from PDGFRα homodimer (top) and PDGFRβ homodimer (bottom) cell lines following a timecourse of PDGF ligand stimulation from 2 min to 4 h with anti-phospho-ERK1/2 (A) or anti-phospho-AKT (C) antibodies. (B, D) Line graphs depicting quantification of band intensities from *n*=3 biological replicates as in A and C. Data are mean ± SEM. Statistical analyses were performed using a ratio paired *t*-test within each cell line and a two-tailed, unpaired *t*-test with Welch’s correction between each cell line. *, p<0.05.

Phosphorylation of AKT in response to PDGFRβ homodimer activation also peaked at 15 min of ligand stimulation (5.852 ± 1.250 R.I.; *p* = 0.0165) but had a greater amplitude than PDGFRα homodimers and appeared more dynamic, sharply declining after the peak at 15 min (**Fig. 6, C and D**). In total, these signaling data suggested that PDGFR homodimers differ in downstream signaling dynamics and that PDGFRβ homodimers generate a greater signaling response to PDGF ligand stimulation than PDGFRα homodimers.

An important question is whether these differences in signaling downstream of the PDGFR homodimers was due to innate differences in the clones that we chose for each of these cell lines, rather than differences in signaling activity downstream of PDGFRα versus PDGFRβ homodimers. We tested this question through activation of an alternate RTK, EGFR. We stimulated each cell line with EGF for 10 min and performed western blotting for the phosphorylation of the same effector molecules, ERK1/2 and AKT, revealing no difference in phosphorylation of these signaling molecules between the PDGFR homodimer cell lines (**Fig. S3, A-D**). These results indicated that the differences that we observed upon PDGF ligand stimulation of our two PDGFR homodimer cell lines reflect true differences in signaling response downstream of these receptors.

### PDGFRβ homodimer activation leads to greater cellular activity

As with many RTKs, signaling through the PDGFRs has been shown to direct a range of cellular processes, such as proliferation, migration and differentiation (Fantauzzo and Soriano, 2015; Schlessinger, 2000). To determine how the above trafficking and signaling dynamics of the PDGFR homodimers affect cellular activity, we first measured proliferation via a soft agar anchorage-independent growth assay in growth media with 10% fetal bovine serum over the course of 10 days. In the absence of ligand, the cells expressing PDGFRβ proliferated more than those expressing PDGFRα, as measured by both colony count (1.1 x 10^4^ ± 1.2 x 10^3^ versus 7.4 x 10^3^ ± 8.6 x 10^2^ colonies; *p* = 0.01) and colony area (7.8 x 10^5^ ± 4.7 x 10^4^ versus 3.3 x 10^5^ ± 4.8 x 10^4^ A.U.; *p* = 0.0002; **Fig. 7, A, B, C, D**). Upon exogenous PDGF ligand stimulation, proliferation as measured by colony count increased for both PDGFRα homodimer (8.6 x 10^3^ ± 8.8 x 10^2^ colonies; *p* = 0.0005) and PDGFRβ homodimer cell lines (1.1 x 10^4^ ± 1.1 x 10^3^ colonies; *p* = 0.01; **Fig. 7, A’, B’, C**). Additionally, the difference in proliferation between PDGFR homodimer cell lines was retained, with the PDGFRβ homodimer cell line exhibiting a higher colony count (*p* = 0.02; **Fig. 7, A’, B’, C**). Exogenous PDGF ligand stimulation did not increase colony area for the PDGFRα homodimer cell line, while it did increase colony area for the PDGFRβ homodimer cell line (8.9 x 10^5^ ± 4.4 x 10^4^ A.U.; *p* = 0.005), again resulting in more proliferation for the PDGFRβ homodimer cell line than the PDGFRα homodimer cell line (*p* = 0.0007; **Fig. 7, A’, B’, D**).

**Figure 7.**
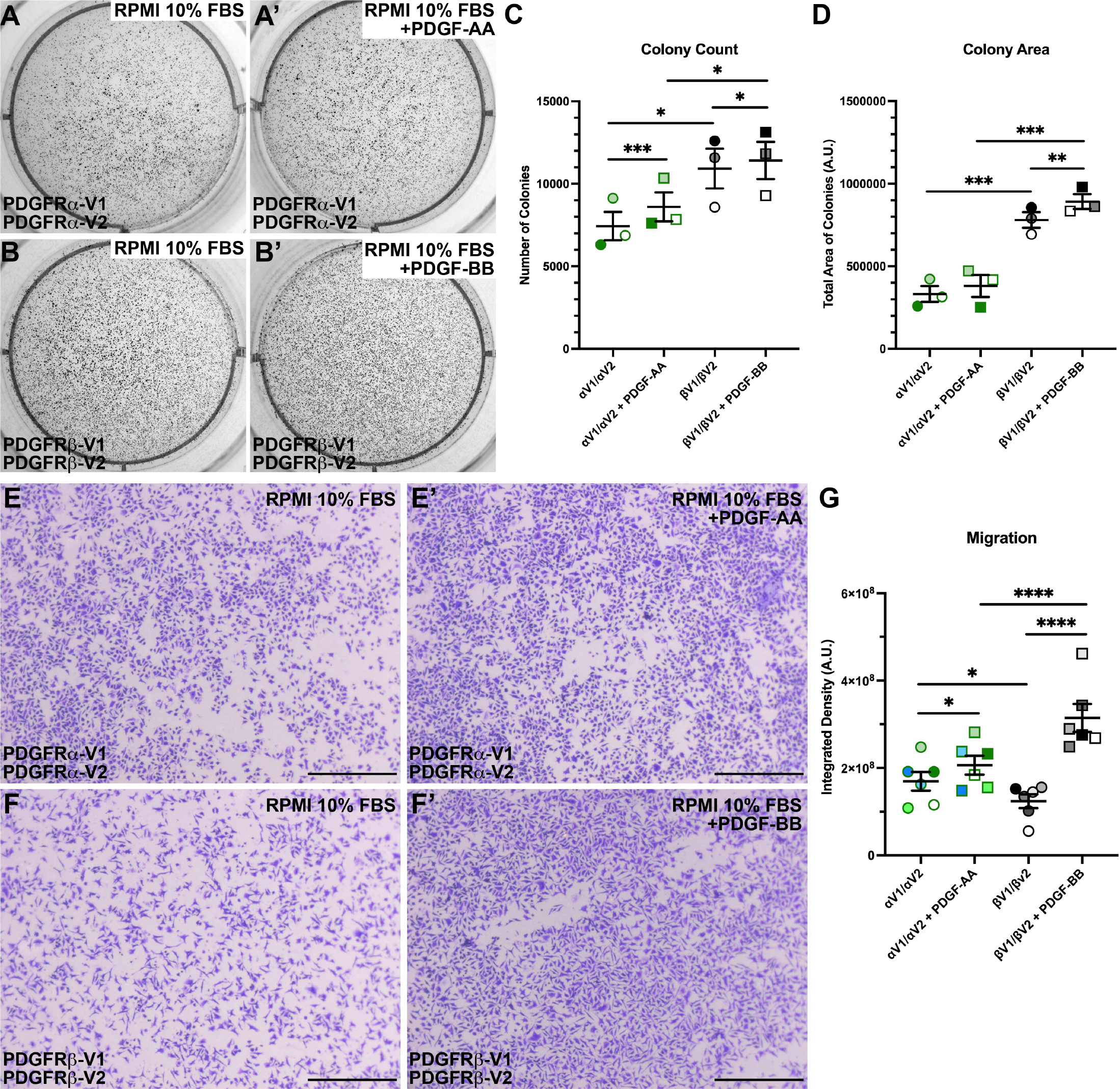
PDGFRβ homodimer activation leads to greater cellular activity. (A-B’) Colony growth in soft agar independent growth assays for PDGFRα homodimer (A-A’) and PDGFRβ homodimer (B-B’) cell lines after 10 days in RPMI growth media in the absence (A, B) or presence (A’, B’) of PDGF ligand. (C-D) Scatter dot plots depicting quantification of colony count (C) or colony area (D) from *n*=3 biological replicates as in A-B’. Data are mean ± SEM. Statistical analyses were performed using a two-tailed, unpaired *t*-test with Welch’s correction between each cell line. p<0.05; **, p<0.01; ***, p<0.001. Colored circles correspond to independent experiments. (E-F’) Crystal violet staining of PDGFRα homodimer (E, E’) and PDGFRβ homodimer (F, F’) cell lines following 24 h of migration through a porous membrane towards RPMI growth media lacking (E, F) or containing (E’, F’) PDGF ligand. Scale bars, 500 µm. (G) Scatter dot plot depicting integrated density in arbitrary units (A.U.) from *n*=6 biological replicates as in E-F’. Data are mean ± SEM. Statistical analyses were performed using a two-tailed, unpaired *t*-test with Welch’s correction between each cell line. p<0.05; ****, p<0.0001. Colored circles correspond to independent experiments.

As a second measure of cell activity, we performed 24 h transwell migration assays in growth media with 10% fetal bovine serum to assess cell migration for the two PDGFR homodimer cell lines. In the absence of exogenous PDGF ligand stimulation, the cells expressing PDGFRα migrated more than those expressing PDGFRβ (1.7 x 10^8^ ± 2.1 x 10^7^ versus 1.2 x 10^8^ ± 1.6 x 10^7^ A.U.; *p* = 0.02; **Fig. 7, E, F, G**). However, while both PDGFRα (2.1 x 10^8^ ± 2.1 x 10^7^ A.U.; *p* < 0.05) and PDGFRβ homodimer cell lines (3.1 x 10^8^ ± 3.2 x 10^7^ A.U.; *p* < 0.0001) exhibited increased migration towards growth media containing PDGF ligand, addition of exogenous ligand resulted in increased migration for cells expressing PDGFRβ over those expressing PDGFRα (*p* < 0.0001; **Fig. 7** **E’, F’, G**). Collectively, these cellular activity assays provided evidence that activated PDGFRβ homodimers generate stronger pro-proliferation and -migration signals than PDGFRα homodimers.

## Discussion

It has previously been impossible to investigate PDGFR dimer-specific dynamics. Former studies have utilized antibody- and ligand propensity-based approaches to study PDGFR expression and/or activity, yet both approaches have considerable caveats. For example, antibodies cannot distinguish whether a receptor is present as a monomer or engaged in a homodimeric or heterodimeric complex. Proximity ligation techniques are also insufficient, as they detect protein-protein interactions within a relatively large range of 40 nm (Bagchi et al., 2015), while dimerized PDGFRs may have considerably closer interactions (Chen et al., 2015). Moreover, evidence suggests that some PDGF ligands are promiscuous and can result in the formation of multiple PDGFR dimers (Fantauzzo and Soriano, 2016). Therefore, in studies where conclusions were drawn about a particular PDGFR dimer based on ligand treatment alone, it is often unclear which dimer was assayed. Thus, our aim was to implement an optimized method, BiFC, to both visualize and purify individual PDGFR dimers to more precisely investigate homodimer-specific subcellular distribution and signaling dynamics. We observed less than complete co-localization between Venus and PDGFR antibody expression, demonstrating that PDGFR expression is not predictive of receptor dimerization and activation, and further indicating that the presence of the BiFC fragments alone does not drive dimerization in our cell lines. The PDGFR-BiFC stable cell lines are thus a viable tool to gain novel insight into dimer-specific activation, trafficking and downstream signaling dynamics.

Our GFP-Trap purification of the PDGFR homodimers revealed increased dimerization for both upon PDGF ligand stimulation, as well as, surprisingly, the presence of dimerized receptors in the absence of ligand stimulation. As recently discussed, an *in silico* study has implicated that there may be an inactive dimerization state for the PDGFRs (Polyansky et al., 2019; Rogers and Fantauzzo, 2020). Interestingly, other RTKs, most notably EGFR, have been shown to exist in an inactive dimerized state in the membrane prior to a ligand-induced conformational change (Bae and Schlessinger, 2010; Zhang et al., 2006). Importantly, even though we detected dimerized PDGFRs in the absence of ligand stimulation, we revealed that PDGFR homodimers were only activated upon exogenous ligand treatment.

In studying these receptor dimerization and activation dynamics, we observed that PDGFRβ receptors dimerized more quickly than PDGFRα receptors. On the other hand, PDGFRα homodimers reached their peak autophosphorylation faster than PDGFRβ homodimers, though the PDGFRα peak was significantly lower than that for PDGFRβ. Based on current evidence in the field, there are two non-mutually-exclusive hypotheses to explain these findings. It is possible that the trends in dimerization and activation observed in our experiments may be due to different conformational changes in the receptors prior to and during activation. Relatedly, it has been shown that ERBB family dimers exhibit both different conformations and varied activation kinetics based on the identity of the activating ligand (Freed et al., 2017). An alternative explanation for the differences in autophosphorylation activity between the PDGFR homodimers may be the auto-inhibitory function of a glutamic acid/proline repeat motif in the C-terminal tail of PDGFRβ (Chiara et al., 2004), which is alleviated upon ligand-induced dimerization and conformational change of the receptor. As this motif has not been detected in PDGFRα, it could create a relative delay in activation for PDGFRβ compared to PDGFRα homodimers.

Our PDGFR homodimer dimerization and activation findings are noteworthy in the context of the Na^+^/K^+^-ATPase and Rab5 co-localization data, from which we determined that PDGFRα homodimers are trafficked from the cell membrane more quickly than PDGFRβ homodimers. Additionally, as discussed above, we demonstrated that PDGFRβ homodimers are activated to a greater extent than PDGFRα homodimers. It may be the case that the extended time at the cell membrane for the PDGFRβ homodimers allows for increased autophosphorylation. Conversely, earlier peak activation of the PDGFRα homodimers may result in faster recruitment of trafficking machinery and, thus, quicker intracellular trafficking.

In addition to differences in the timing of initial PDGFR trafficking into early endosomes, our data also revealed varied degradation dynamics and recycling between the PDGFR dimers. We found that PDGFRα homodimers co-localized more with Rab7 following extended PDGF ligand stimulation and were degraded faster than PDGFRβ homodimers. Interestingly, however, both PDGFR homodimers experienced similar levels of receptor degradation following 4 h of ligand treatment. We additionally found that PDGFRβ homodimers were more likely to be recycled than PDGFRα homodimers. However, the fact that only 20-22% of initial receptor levels for both cell lines were detected following 4 h of PDGF ligand treatment indicated that receptor degradation is the predominant result of intracellular trafficking, consistent with previous findings for PDGFRβ (Hellberg et al., 2009; Sadowski et al., 2013).

We next sought to determine how these dimerization and activation dynamics, together with PDGFR dimer-specific subcellular distribution, affected downstream signaling. We found that both PDGFR homodimers induced phospho-ERK1/2 peaks that remained relatively stable over time. PDGFRα homodimers drove an early peak of ERK1/2 phosphorylation, while the peak driven by PDGFRβ homodimer activation occurred slightly later and had a higher amplitude. Interestingly, it has previously been shown that various subcellular localizations of Erk1/2, or its upstream activator RAS, result in different timing of the Erk1/2 signaling response (Bruggemann et al., 2021; Herrero et al., 2016; Keyes et al., 2020). For example, following EGF ligand stimulation, the phosphorylation of Erk1/2 at the membrane results in sustained signaling, while its phosphorylation in the cytoplasm and nucleus results in transient signaling, ultimately leading to differences in cellular activity (Keyes et al., 2020). Given that PDGFRs have been shown to signal within various cellular compartments, including endosomes (Wang et al., 2004), engagement of signaling molecules at different subcellular locations may explain, at least in part, the observed differences in ERK1/2 signaling dynamics downstream of PDGFRα versus PDGFRβ homodimer activation. Correspondingly, the trends of AKT phosphorylation were also different between the two PDGFR homodimers. Though initial phospho-AKT peaks occurred at a similar timepoint for both PDGFR homodimers, the PDGFRα homodimer-driven peak remained relatively stable, while PDGFRβ homodimer-stimulated phosphorylation of AKT was much more transient. In line with the phospho-ERK1/2 results, PDGFRβ homodimers displayed a greater level of phospho-AKT than PDGFRα homodimers, suggesting that signaling through PDGFRβ homodimers results in greater activation of downstream signaling cascades. Taken together, these findings support a model in which differences in the timing and extent of signaling dynamics downstream of PDGFR activation fine tune downstream biological responses. An important consideration, however, is that we evaluated whole cell lysates reflecting population-level dynamics, and it is likely that these dynamics differ between individual cells (Sparta et al., 2015). Analysis of single-cell signaling dynamics in response to PDGFR dimer-specific activation is beyond the scope of this study, but will be a focus of future work.

Finally, cell activity assays revealed that PDGF ligand stimulation of PDGFRβ homodimers resulted in increased levels of both proliferation and migration from what was observed upon PDGFRα homodimer activation. Interestingly, PDGFRα homodimers induced more migration than PDGFRβ homodimers towards growth media alone, which contains PDGF ligands in the serum, while the addition of exogenous PDGF ligand led to a more significant increase in migration for the PDGFRβ homodimer cell line. It is possible that the low level of PDGF ligand present in growth media is stimulatory towards PDGFRα homodimer-driven migration, but that higher levels of ligand are required for PDGFRβ homodimer-driven migration. Along these lines, varying concentrations of PDGF ligand have been shown to result in differences in both PDGFR trafficking and downstream migration and proliferation (De Donatis et al., 2008). These varying responses to changes in ligand concentration could serve as a mechanism to introduce biological specificity downstream of PDGFR dimerization. Alternatively, activation of PDGFRβ homodimers may generate stronger kinase activity than that of PDGFRα homodimers. In fact, analysis of chimeric PDGFRs *in vivo* revealed that the intracellular domain of PDGFRβ can compensate for the loss of PDGFRα signaling, while the inverse is not true (Klinghoffer et al., 2001). Both hypotheses considered, it remains that PDGFRβ homodimers stimulated increased cell activity upon exogenous PDGF ligand stimulation in the context of HCC15 cells.

Overall, we have identified PDGFR homodimer-specific differences in dimerization, activation, trafficking and downstream signaling and cellular activity utilizing a novel BiFC system. Our findings thus provide significant insight into how similar receptors within an RTK family utilize a combination of mechanisms to differentially propagate downstream signaling. This approach will be invaluable in future studies characterizing PDGFR dimer-specific interactomes to further delineate the means by which these receptors generate distinct cellular outputs.

## Abbreviations

EGFR: epidermal growth factor receptor
BiFC: bimolecular fluorescence complementation
PCC: Pearson’s correlation coefficient
PDGFR: platelet-derived growth factor receptor
R.I.: relative induction
RTK: receptor tyrosine kinase

## Acknowledgements

We thank Robert Long, Julia Mo, Damian Garno and Jessica Patrick for technical assistance. Cell sorting was performed at the University of Colorado Cancer Center Flow Cytometry Shared Resource (Cancer Center Support Grant P30CA046934). We are grateful to the lab of Dr. Rytis Prekeris for pLVX-puro, pCMV-VSV-G and pCMV-dR8.91 plasmids and the lab of Dr. Lynn Heasley for HCC15 cells. We thank members of the Fantauzzo laboratory and Dr. Rytis Prekeris for their critical comments on the manuscript. This work was supported by the National Institutes of Health (R01DE027689 to K.A.F., K02DE028572 to K.A.F. and F31DE029976 to M.A.R.). The authors declare no competing financial interests.

## Author contributions

Conceptualization: K.A. Fantauzzo and M.A. Rogers; Methodology: M.A. Rogers, K.A. Fantauzzo; Formal analysis: M.A. Rogers; Investigation: M.A. Rogers, K.A. Fantauzzo; Writing – original draft: M.A. Rogers; Writing – review & editing: M.A. Rogers, K.A. Fantauzzo; Visualization: M.A. Rogers, K.A. Fantauzzo; Supervision: K.A. Fantauzzo; Funding acquisition: M.A. Rogers, K.A. Fantauzzo

## Notes

### Competing Interest Statement

The authors have declared no competing interest.

