## Supplemental Material for "PDGFR dimer-specific activation, trafficking and downstream signaling dynamics"

##### Supplemental figure legends

**Figure S1. Validation of PDGFR-BiFC stable cell lines.** (A) Schematic of PDGFRα fused to the non-fluorescent N-terminal (V1) fragment of Venus and PDGFRα fused to the C-terminal (V2) fragment. Upon receptor dimerization, a functional Venus protein is generated. (B-C') Venus expression (white/green) as assessed by fluorescence analysis of nontransduced HCC15 cells in the absence (B, B') or presence (C, C') of PDGF ligand for 5 min. Nuclei were stained with DAPI (blue; B', C'). *n*>20 technical replicates across each of 2 biological replicates. Scale bars, 20 µm. (D-E) Western blot (WB) analysis of whole cell lysates (WCL) from nontransduced (NT) HCC15 cells (left), PDGFRα homodimer (D; right) and PDGFRβ homodimer (E; right) cell lines in the absence or presence of PDGF ligand for 15 min with anti-PDGFRα (top) or anti-PDGFRβ (bottom) antibodies. *n*=2 biological replicates per condition.

**Figure S2. PDGFRβ homodimers are more likely to localize to the cell membrane following 90 minutes of PDGF ligand stimulation than PDGFRα homodimers.** (A) Bar graph depicting Pearson's correlation coefficient of the PDGFRα homodimer and PDGFRβ homodimer cell lines with an anti-$Na^+/K^+$-ATPase antibody following PDGF stimulation for 90 min. Data are mean ± SEM. Statistical analyses were performed using a two-tailed unpaired *t*-test with Welch's correction. *, $p<0.05$. Colored circles correspond to independent experiments. *n*>20 technical replicates across each of 2 biological replicates. (B-C") $Na^+/K^+$-ATPase antibody expression (white/magenta; B, B",

C, C”) and/or Venus expression (white/green; B’, B”, C’, C”) as assessed by (immuno)fluorescence analysis of PDGFRα homodimer (B-B”) and PDGFRβ homodimer (C-C”) cell lines. Insets in B” and C” are regions where white arrows are pointing. Nuclei were stained with DAPI (blue; B”, C”). White arrows denote co-localization; white outlined arrows denote lack of co-localization. Scale bars, 20 μm. Inset scale bars, 3 μm.

**Figure S3. Phosphorylation of ERK1/2 and AKT downstream of EGF stimulation is unchanged between stable PDGFR-BiFC cell lines.** (A, C) Western blot (WB) analysis of whole cell lysates (WCL) from PDGFRα homodimer (left) and PDGFRβ homodimer (right) cell lines in the absence of ligand or following EGF ligand stimulation for 10 min with anti-phospho-ERK1/2 (A) or anti-phospho-AKT (C) antibodies. (B, D) Scatter dot plots depicting quantification of band intensities from $n$=3 biological replicates as in A and C. Data are mean ± SEM. Statistical analyses were performed using a two-tailed, unpaired $t$-test with Welch’s correction between each cell line. ns, not significant. Shaded circles correspond to independent experiments.

### Figure S1.

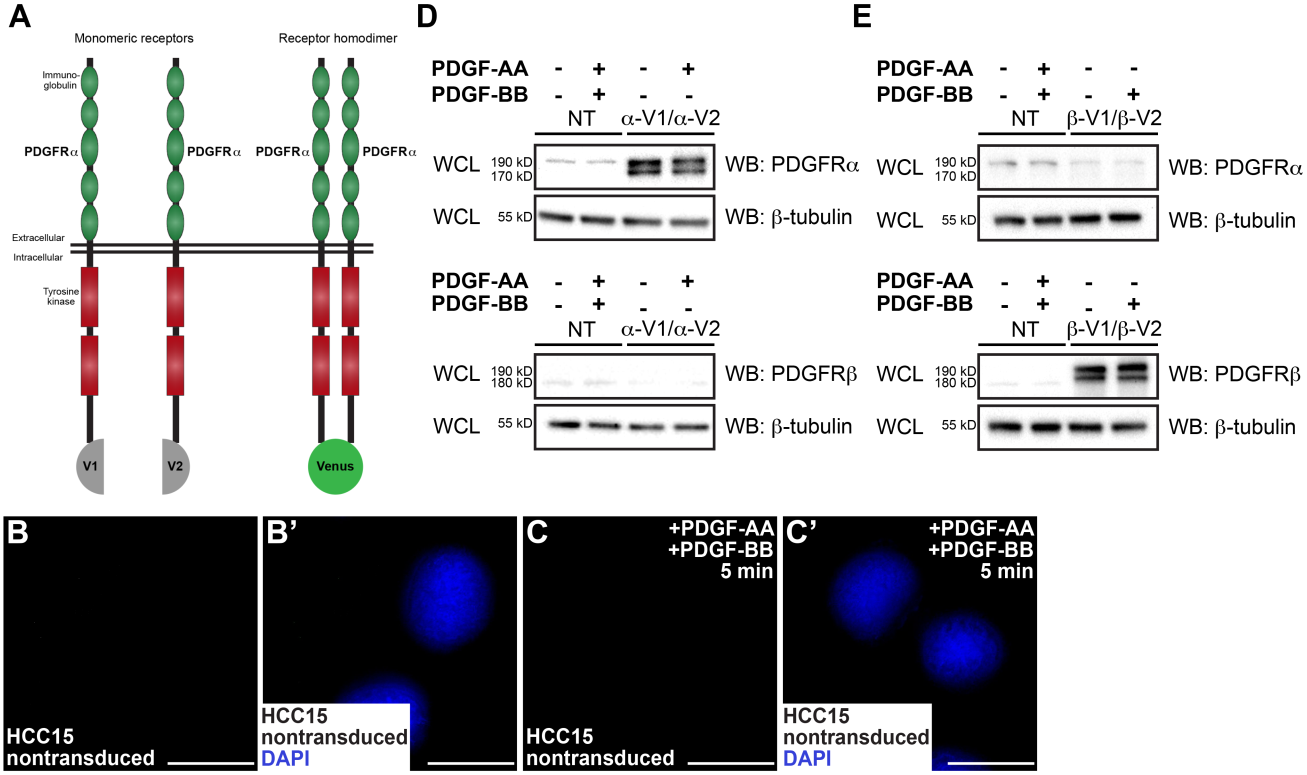
A
Monomeric receptors
Receptor homodimer
Immuno-globulin
PDGFRα
PDGFRα
PDGFRα
PDGFRα
Extracellular
Intracellular
Tyrosine kinase
V1
V2
Venus
D
PDGF-AA - + - +
PDGF-BB - + - -
NT
α-V1/α-V2
WCL 190 kD 170 kD WB: PDGFRα
WCL 55 kD WB: β-tubulin
PDGF-AA - + - +
PDGF-BB - + - -
NT
α-V1/α-V2
WCL 190 kD 180 kD WB: PDGFRβ
WCL 55 kD WB: β-tubulin
E
PDGF-AA - + - -
PDGF-BB - + - +
NT
β-V1/β-V2
WCL 190 kD 170 kD WB: PDGFRα
WCL 55 kD WB: β-tubulin
PDGF-AA - + - -
PDGF-BB - + - +
NT
β-V1/β-V2
WCL 190 kD 180 kD WB: PDGFRβ
WCL 55 kD WB: β-tubulin
B
HCC15 nontransduced
B'
HCC15 nontransduced DAPI
C
+PDGF-AA +PDGF-BB 5 min
HCC15 nontransduced
C'
+PDGF-AA +PDGF-BB 5 min
HCC15 nontransduced DAPI

### Figure S2.

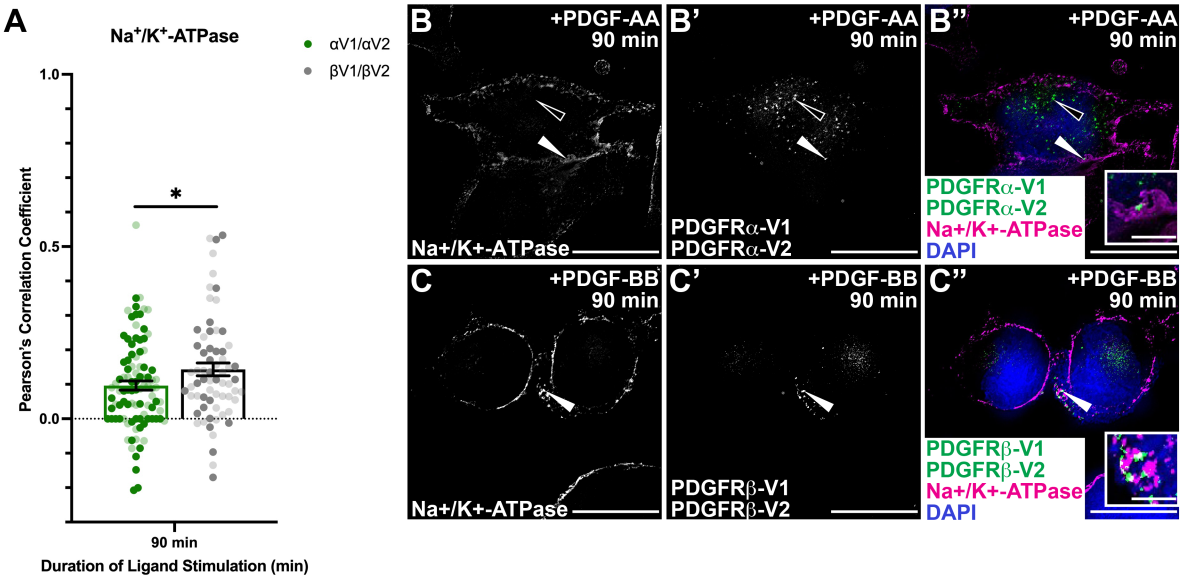
A
Na+/K+-ATPase
αV1/αV2
βV1/βV2
1.0
0.5
0.0
*
Pearson's Correlation Coefficient
90 min
Duration of Ligand Stimulation (min)
B
+PDGF-AA
90 min
Na+/K+-ATPase
B'
+PDGF-AA
90 min
PDGFRα-V1
PDGFRα-V2
B''
+PDGF-AA
90 min
PDGFRα-V1
PDGFRα-V2
Na+/K+-ATPase
DAPI
C
+PDGF-BB
90 min
Na+/K+-ATPase
C'
+PDGF-BB
90 min
PDGFRβ-V1
PDGFRβ-V2
C''
+PDGF-BB
90 min
PDGFRβ-V1
PDGFRβ-V2
Na+/K+-ATPase
DAPI

#### Figure S3.

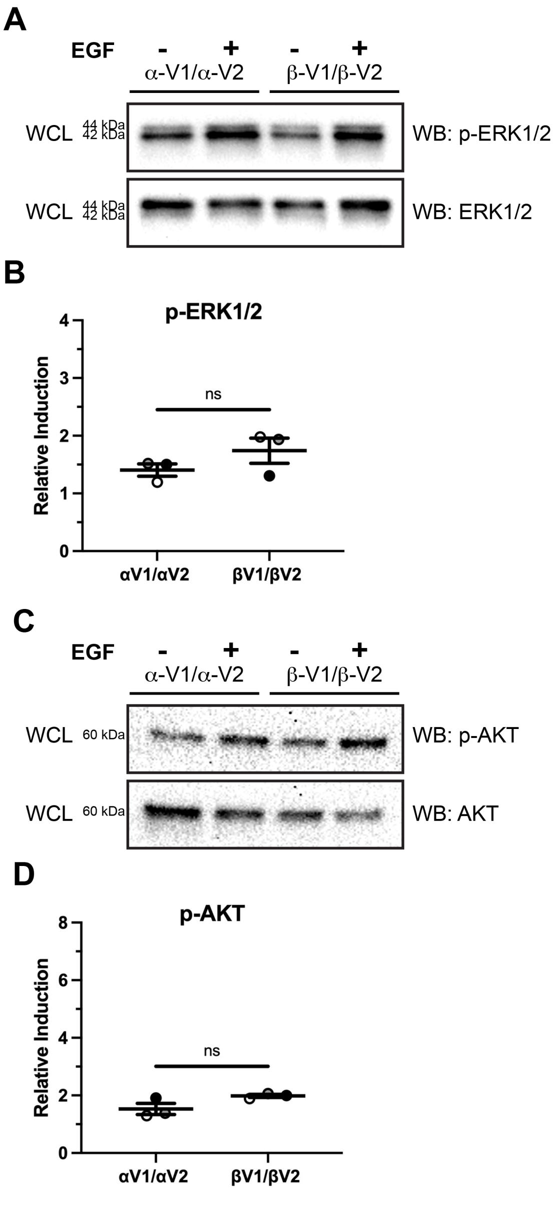

A
EGF
-
+
-
+
α-V1/α-V2
β-V1/β-V2
WCL
44 kDa
42 kDa
WB: p-ERK1/2
WCL
44 kDa
42 kDa
WB: ERK1/2
B
p-ERK1/2
Relative Induction
ns
4
3
2
1
0
αV1/αV2
βV1/βV2
C
EGF
-
+
-
+
α-V1/α-V2
β-V1/β-V2
WCL
60 kDa
WB: p-AKT
WCL
60 kDa
WB: AKT
D
p-AKT
Relative Induction
ns
8
6
4
2
0
αV1/αV2
βV1/βV2
